# A citywide metagenomic analysis reveals surface-specific microbiome and resistome patterns in outdoor urban environments across Liverpool, UK

**DOI:** 10.64898/2026.02.18.706585

**Authors:** Gavin Ackers-Johnson, Angus M. O’Ferrall, Andrew Holmes, Ellie Allman, Pauline Ambrose, Anya Breen, Amber Cutcliffe, Kara D’Arcy, Richard Goodman, Alexander Kingdon, Amy McLeman, Sabrina Moyo, Ralfh Pulmones, Moon Deb Rajib, Priyanka Sharma, Ellinor Shore, Karina Clerkin, Maria Moore, Helen McNeil, Adam P. Roberts

## Abstract

Urbanisation is rapidly increasing worldwide, with increasing attention focused on its consequences for human populations. Despite the importance of outdoor urban environments for biodiversity and human wellbeing, their microbial ecology remains poorly characterised, particularly in relation to emerging microbial threats including antimicrobial resistance (AMR). Where previous studies have targeted individual urban niches, we used shotgun metagenomic sequencing to assess microbial community composition and antimicrobial resistance genes (ARGs) across five outdoor surface types at a citywide level: Touchpoints, Pathways, Soil, Refuse and Watersides.

Microbial community composition differed significantly between surface types, with surface identity explaining 33.1% of variation, compared with 15.7% explained by council ward. Soil harboured the most compositionally distinct communities, while Touchpoints and Refuse showed greater overlap, with Pathways and Watersides occupying an intermediate position between the Touchpoint/Refuse and Soil communities. Touchpoints had significantly higher ARG abundance than Pathways, Soil and Watersides, with Refuse showing greater variability driven largely by a single high-abundance observation; following sensitivity analysis, Touchpoints were also significantly higher than Refuse. ARGs were rarely detected in Pathways, Watersides and Soil.

These findings provide a citywide baseline showing that outdoor urban surfaces support distinct microbial communities and resistance profiles, with the strongest ARG signals observed on frequently contacted surfaces. This highlights the value of surface-specific metagenomic surveillance for characterising the urban microbial and antimicrobial resistance landscape.

**Data Summary:** Trimmed metagenomic sequence reads have been deposited in the NCBI Sequence Read Archive under BioProject accession number PRJNA1424689.

Source code and data for reproducing figures are available at: https://github.com/gaj-bio/Liverpool_microbiome_metagenomics.

## Introduction

The share of the global population residing in urban areas is increasing across all regions of the world, with over 55% of people currently living in such settings (approximately 4.2 billion) and an additional 2.5 billion people projected to do so by 2050. This proportion rises to 81% in high-income countries and 85% in the United Kingdom [1]. These urban environments represent highly complex ecosystems where human activity, infrastructure, and natural elements converge to shape microbial communities. Alongside a growing built environment to accommodate high-density populations, including housing, healthcare and transport, outdoor environments also serve as hubs of human activity, with exposure to urban green spaces increasingly associated with a range of beneficial health-related outcomes [2, 3]. A diverse array of microorganisms are able to colonise these unique niches, collectively influencing public health, environmental resilience and microbial evolution through broadly variable and poorly understood mechanisms [4].

A growing interest has been placed on the built environment and indoor microbiome as having a profound influence on human health, with urbanisation associated with heightened risks of depression, obesity, allergic disease, respiratory disease and other conditions. These have been linked to reduced microbial exposure through diminished microbial diversity and increased levels of pollutants and antimicrobial chemicals [5]. The latter intersects directly with the challenge of antimicrobial resistance (AMR), one of the top threats to global public health, with bacterial AMR estimated to be directly responsible for 1.27 million and a contributing factor towards 4.95 million global deaths in 2019 [6].

Anthropogenic pressures are more diffuse in outdoor environments, where temperature, moisture, and nutrient availability fluctuate unpredictably, yet they continue to exert a strong influence on both taxonomic composition and AMR dynamics [7]. Human waste, aerosols, runoff, and microbial immigration continuously introduce microorganisms, antibiotic residues, metals, and other co-selecting agents into outdoor settings. These inputs collectively shape community structure and promote the persistence and potential enrichment of resistance elements [8].

Outdoor environments are also more susceptible to climate change. Alongside general temperature increases and shifts in precipitation patterns, more frequent and intense extreme weather events are predicted [9]. These climatic stressors can alter microbial survival, dispersal, and horizontal gene exchange, reshaping both taxonomic composition and functional potential [10, 11]. Specifically, studies in the USA, Europe and China have shown higher ambient temperatures are associated with a greater prevalence of AMR infections [12–14].

Despite growing interest, characterisation of urban microbiomes has largely been fragmented. Most studies to date have targeted specific environments, either where there is a high risk of infection and ARG transmission (e.g. hospitals [15, 16] and wastewater treatment systems [17, 18]) or areas of the built environment with high human traffic (e.g. indoor spaces [4, 19, 20] and transport systems [21, 22]). Although these studies yield valuable insights into urban microbial dynamics, their scope is often compartmentalised.

Here, we aimed to adopt a holistic, city-wide approach to characterising microbial diversity. We conducted shotgun metagenomic sequencing on 125 samples collected from five distinct types of outdoor urban surface: human touchpoints, pathways, soil, refuse, and waterside environments. These surface types were selected to represent contrasting urban microhabitats shaped by different levels of human contact, environmental exposure, moisture availability, and nutrient inputs. Human touchpoints represent surfaces subject to frequent direct human interaction and associated microbial inputs, while pathways experience high levels of human traffic but greater exposure to environmental stresses such as desiccation and UV radiation. In contrast, soil represents a more heterogeneous environmental reservoir with greater potential for moisture and organic matter retention, while refuse provides a distinct environment characterised by concentrated organic inputs and human-associated material. Waterside environments represent moist interfaces between urban land and water, where water availability and urban runoff may influence microbial persistence and composition. We therefore expected these contrasting microhabitats to support distinct microbial communities and resistome profiles, with differences reflecting the relative influence of human-associated inputs, environmental exposure, substrate characteristics, and moisture and nutrient availability.

We characterised taxonomic composition at the phylum and genus levels, and examined the resistome by quantifying the abundance of ARGs. Through this integrated approach, we provide a city-scale assessment of the outdoor urban microbiome and its resistance potential. This baseline data lays the foundation for future comparative analyses under environmental change and supports ongoing surveillance of environmental AMR as part of a One Health framework.

## Methods

### Sample Collection and Study Design

Twenty-five council wards across Liverpool were sampled, prioritising the city centre and adjacent areas (**Figure 1A**). Ward-associated data are made available to the public by Liverpool City Council [23]. Within each ward, five outdoor surface categories were sampled to capture a broad range of ecological niches: Touchpoints, Pathways, Soil, Refuse, and Waterside. In total, 125 samples were collected (one sample per surface type per ward). All sampling was conducted exclusively in publicly accessible outdoor areas, with no collections taken from private residential or commercial premises.

**Figure 1.**
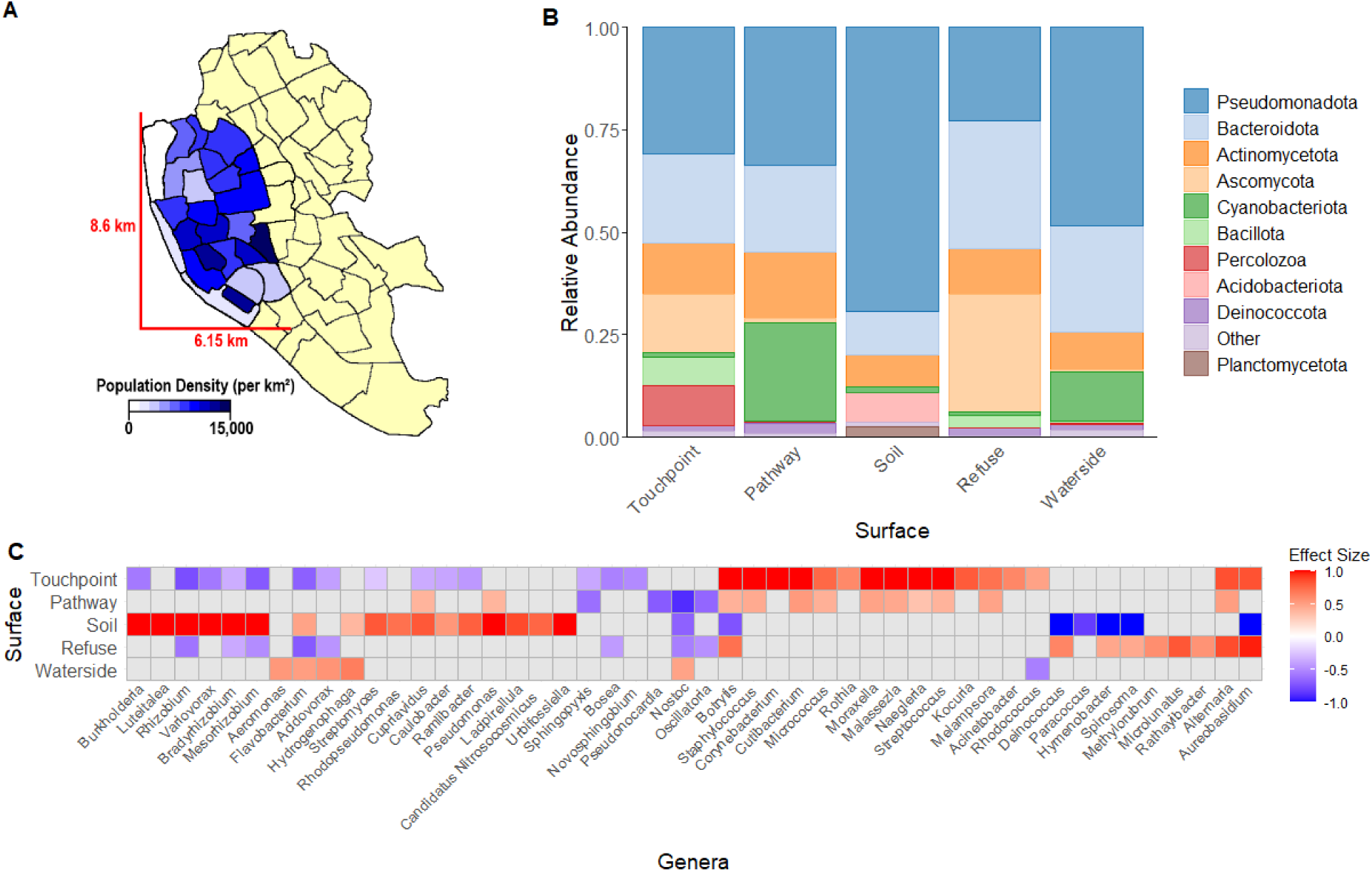
**A)** Map of Liverpool City wards showing population density for the 25 sampled wards (highlighted blue). Wards outside the sampling area are indicated in yellow. **B)** Relative abundance of phyla across surface types. The five most abundant phyla from each surface type were identified and combined, resulting in a total of ten phyla displayed. All remaining phyla are grouped under “Other”. **C)** Differentially abundant genera identified using ALDEx2. A 10% global prevalence filter was applied, and each surface type was compared to the mean of the remaining four surface types. Only genera with a statistically significant difference (p < 0.05) and an effect size exceeding ± 0.5 in at least one surface type are shown. Genera are clustered based on effect size across surface types, with effect size indicated by a red (positive) to blue (negative) colour gradient.

Touchpoints were defined as surfaces explicitly designed for frequent human hand interaction, including pedestrian crossing buttons, letterboxes, and playground equipment. Pathways included surfaces subject to regular foot traffic such as pavements and footpaths. Soil samples were collected from urban nature interfaces, including flowerbeds, tree bases, and grassed areas. Refuse surfaces comprised bins, litter, and fly-tipped materials associated with waste disposal. Waterside samples were taken adjacent to bodies of water (including ponds, lakes, canals, and temporary water features such as puddles) by swabbing the solid surface at the water–ground interface.

Sampling took place over two consecutive days (26^th^ and 27^th^ September 2024) between 10:00 and 14:00. During sampling, temperatures ranged from approximately 12 °C to 15 °C on both days. On day one, sampling began during light to moderate rainfall, with overcast conditions persisting for the remainder of the sampling period. Day two remained dry, with high cloud cover during the early sampling period that gradually cleared throughout the day. Weather conditions during the week preceding sampling were broadly similar, being predominantly dry and overcast with intermittent light to moderate rainfall.

For Soil samples, surface soil from a depth of up to 2 cm was collected directly into 15 mL Falcon tubes using the 3 mL mark as a guide; in the laboratory, 200 mg of soil was subsequently transferred into 1 mL DNA/RNA Shield buffer. All other surfaces were sampled using Zymo Research DNA/RNA Shield collection swabs (1 mL; catalogue R1106). Each respective surface was vigorously swabbed back and forth multiple times across the accessible surface to recover as much microbial biomass as possible. An area of approximately 25 cm² was targeted where practicable, although the precise area could not be standardised across samples because of the heterogeneity of the surfaces assessed. Two low-biomass contamination controls were used: one blank Zymo Research DNA/RNA Shield collection swab and one blank 1 mL DNA/RNA Shield buffer tube. Both controls were processed alongside the environmental samples through DNA extraction and library preparation.

### Sample Locations

Sample geolocations and surface metadata are available in **Supplementary Material 1** and online (https://www.google.com/maps/d/edit?mid=1jfQz8wkxY-bCkOD6yKjqpL1fH1fBWw0&usp=sharing).

### DNA Extraction, Library Preparation and Sequencing

Samples preserved in DNA/RNA Shield were shipped at ambient temperature to MicrobesNG (Birmingham, UK) for DNA extraction, Illumina library preparation, and shotgun metagenomic sequencing. The DNA concentration extracted from each sample and its respective concentration post-library preparation are available in **Supplementary Material 2**. The target input used for the library preparation was 0.5 ng/µL, with DNA concentrations below this threshold used at their available concentration. Libraries were prepared using the Illumina Nextera XT kit with modified input and PCR conditions. Libraries were sequenced using 2 × 250 bp paired-end reads with a target yield of ≥2 million reads per sample. Reads were quality-trimmed using Trimmomatic v0.39 [24] and assessed using fastp v0.23.2 [25] and internal MicrobesNG QC tools. Samples with <500,000 read pairs or with an overlapping insert-size peak deviating from ∼400 bp were removed prior to downstream analysis.

### Taxonomic Profiling

Taxonomic classification was performed by MicrobesNG (Birmingham, UK) using KrakenUniq v0.6 [26], retaining read-based taxonomic assignments passing a >0.02% unique k-mer threshold under the sensitive filter. This resulted in the identification of 490 taxa at the genus level across bacteria and fungi following application of the >0.02% unique k-mer threshold.

Abundance re-estimation was conducted by MicrobesNG (Birmingham, UK) using Bracken v2.7 [27], and we used the genus-level Bracken sensitive reports as the primary read-count input for downstream taxonomic quantification and diversity analyses. NCBI Taxonomy IDs were used to aggregate counts to higher ranks (including family and phylum). Samples and metadata were imported into R v4.5.1 and structured into a Tree Summarized Experiment object with the mia v1.16.0 package for community profiling [28].

The top five most abundant phyla or genera on each surface type were identified, with all others designated “Other”, and visualised with the plotAbundance function of miaViz v1.16.0. Phylum-level differences among surface types were tested using Kruskal-Wallis rank-sum tests followed by Dunn’s post-hoc comparisons with Benjamini-Hochberg FDR correction using the FSA v0.10.0 package [29].

Differential abundance analysis of genus-level taxa was performed using ALDEx2 v1.40.0 to account for the compositional nature of metagenomic read counts [30]. Bracken-derived genus counts were first filtered to retain genera present in at least 10% of samples to reduce sparsity. Counts were then transformed using ALDEx2’s CLR approach, with Monte Carlo sampling (N = 128) from the Dirichlet distribution to estimate within-condition technical variation. Differential abundance was evaluated by comparing each surface type against the mean of all remaining surfaces (i.e., one-vs-rest contrasts). Significant genera were defined as those with Benjamini–Hochberg adjusted p-values < 0.05 and an effect size magnitude > 0.5. Genera meeting the thresholds were visualised using an effect-size heatmap to highlight surface-associated taxa.

### Diversity and Community Structure Analysis

Alpha diversity metrics (Observed richness, Shannon diversity, Faith’s Phylogenetic Diversity) were calculated at the genus level from Bracken-derived taxonomic count data using the ‘addAlpha()’ function within the Tree Summarized Experiment/mia framework in R [28]. Diversity estimates were generated directly from non-rarefied count data to retain the available sequencing information and avoid discarding reads through subsampling. Statistical differences in alpha diversity among surface types were tested using pairwise Wilcoxon rank-sum tests, with Benjamini–Hochberg correction applied for multiple comparisons (FDR < 0.05). Only significant pairwise differences were displayed on plots.

Beta diversity was estimated at the genus level using Bray-Curtis dissimilarity calculated from relative abundance data within the Tree Summarized Experiment/mia framework in R [28]. Ordination was performed using both metric multidimensional scaling (MDS) and non-metric multidimensional scaling (NMDS) implemented using ‘runMDS()’ and ‘runNMDS()’, respectively. MDS eigenvalues were extracted and converted to proportional variance contributions for axis interpretation, while NMDS ordination included automatic computation of stress, which was used to assess goodness-of-fit.

To evaluate the influence of environmental factors on community structure, distance-based redundancy analysis (dbRDA) was performed using Bray-Curtis distances with ‘addRDA()’, testing the effects of Surface and Council Ward. Significance of community differences was assessed using PERMANOVA (‘getPERMANOVA()’) applied to Bray-Curtis matrices, with p-values obtained from permutation-based tests. Variance explained by each factor was reported as R². Tests for homogeneity of multivariate dispersions were implemented within the mia workflow to evaluate whether observed group differences might reflect heterogeneity in dispersion rather than centroid separation.

### Antimicrobial Resistance Gene Detection

Antimicrobial resistance gene (ARG) detection was performed using a read-based alignment workflow. The ResFinder reference fasta (v2.3.2) was indexed prior to alignment, and quality-trimmed paired-end reads (MicrobesNG output) were mapped directly to the indexed ResFinder database using k-mer alignment via KMA [31]. Raw ‘.res’ files were then filtered to retain high confidence matches using thresholds of >1 read, ≥90% nucleotide identity, and ≥60% gene coverage. For exploratory analyses, a relaxed filter set was applied using the same parameters but with ≥30% gene coverage to enable detection of lower-abundance resistance genes. ResFinder was selected due to its well-curated, regularly updated database of acquired resistance genes and its widespread use in metagenomic and genomic surveillance studies, enabling standardised and comparable ARG detection across datasets.

Filtered ARG hits were normalised using reads per kilobase per million reads (RPKM) to account for gene length and sequencing depth. The summed RPKM values across ARGs were used as a measure of overall sequencing-derived ARG abundance per sample. Differences in overall ARG abundance across surfaces were assessed using pairwise Wilcoxon rank-sum tests with Benjamini-Hochberg FDR correction. ARGs were then grouped by antimicrobial class, and relative class contributions were visualised using stacked bar plots to compare resistome profiles across surface types.

To examine gene-level resistome structure, ARG abundance tables were converted to a binary presence/absence matrix, where detections were coded as 1 and non-detections as 0. Because ARGs were mainly identified in Touchpoint and Refuse environments, downstream comparison focused on these two surface types. Genes detected in at least one sample from either category were retained, and binary heatmaps were generated to visualise shared and unique ARGs across surfaces, with ARGs grouped by antimicrobial class.

## Results

### City-wide metagenomic profiling reveals distinct and shared microbial taxa across outdoor surfaces

All samples (n = 125) underwent shotgun metagenomic sequencing, generating a median of 11.59–12.52 million paired-end reads per surface type, except for Touchpoints, which yielded 15.72 million (**Supplementary Figure 1A**). This may reflect differences in microbial biomass and/or DNA yield between surface types. The median number of reads taxonomically classified using KrakenUniq ranged from 1.53 to 1.65 million for most surface types; however, Waterside and Soil samples produced lower values of 1.06 and 0.65 million respectively (**Supplementary Figure 1B**). The distribution of genera exhibited a characteristic long-tailed pattern, where the majority of taxa were low in both abundance and prevalence, while a small subset dominated in both metrics. This trend was consistently observed across all surface types (**Supplementary Figure 2**).

At the phylum level, Pseudomonadota was the most abundant taxon (**Figure 1B; Supplementary Material 3**), with relative abundances ranging from 23% to 69% across surface types. Its highest abundance occurred in Soil (69%), which was significantly greater than in all other surfaces (Kruskal–Wallis [KW] test, p < 0.001; Dunn’s post hoc tests with Benjamini–Hochberg correction, all adjusted p < 0.005). Refuse was the only surface where Pseudomonadota did not predominate (23%); instead, Bacteroidota (31%) and Ascomycota (28%) were most abundant.

While Bacteroidota was relatively abundant across all surfaces (11–31%), Ascomycota showed a more restricted distribution, being abundant only in Refuse (29%) and Touchpoints (14%), where its abundance was significantly higher than in all other surfaces (KW test, p < 0.001; Dunn’s test, BH-adjusted p < 0.001). Conversely, Cyanobacteriota was abundant primarily on Pathways (24%) and Watersides (12%), again at significantly higher levels than on other surfaces (KW test, p < 0.001; Dunn’s test, BH-adjusted p < 0.005). Finally, the abundances of Acidobacteriota and Percolozoa were significantly higher in Soil (7%) and Touchpoints (10%), respectively, compared with all other surface types (KW test, p < 0.001; Dunn’s test, BH-adjusted p < 0.001).

To assess potential heterogeneity within the surface categories, individual sample phylum-level abundance profiles were examined, with samples ordered according to Bray–Curtis dissimilarities (**Supplementary Figure 3**) and associated sampling metadata groupings (**Supplementary Figure 4**). These profiles demonstrated clear within-surface variability, while also showing broad patterns consistent with the overall differences observed between surfaces. No individual samples or specific surface subtypes showed a clear, distinct abundance profile that would provide an objective basis for exclusion or indicate that the observed surface-level patterns were driven by a particular sample or subtype. Consequently, no samples or subtypes were excluded on this basis.

The relative abundance profiles of the most common genera further highlighted the distinctiveness of the Soil microbiome, which showed the broadest diversity and the most differentiated abundance patterns across surface types (**Supplementary Figure 5**). To statistically assess these patterns, we performed a genera-level differential abundance analysis using ALDEx2, contrasting each surface against the mean of all other surfaces and visualising those with a statistically significant difference (p < 0.05) and an effect size exceeding ± 0.5 in at least one surface type (**Figure 1C**). Genera were pre-filtered by a ≥10% prevalence threshold to avoid differential abundance estimates driven by sparsely detected taxa. This contrast-based approach revealed that Soil and Touchpoints displayed the strongest surface-specific signatures, with the largest positive and negative effect sizes across genera. When averaged across the filtered genera set, Soil and Touchpoints showed mean effect sizes of 0.66 and 0.52, respectively – substantially higher than those observed for Pathways (0.30), Refuse (0.32), and Watersides (0.23).

Only a subset of genera exhibited effect sizes exceeding ±1, corresponding to more than two-fold shifts in centred log-ratio (CLR) space. In Touchpoints, enriched genera included *Botrytis* (1.00), *Streptococcus* (1.02), *Moraxella* (1.11), *Malassezia* (1.14), *Naegleria* (1.17), *Corynebacterium* (1.28), *Staphylococcus* (1.39), and *Cutibacterium* (1.45). Soil showed an even broader set of enriched genera, including *Variovorax* (1.06), *Pseudomonas* (1.12), *Urbifossiella* (1.20), *Mesorhizobium* (1.22), *Rhizobium* (1.38), *Bradyrhizobium* (1.39), and *Burkholderia* (1.85). The only genus with an effect size exceeding 2 was *Luteitalea* (2.60), also in Soil. Large negative effect sizes (≤ −1) were found exclusively in Soil, including *Spirosoma* (−1.12), *Aureobasidium* (−1.16), *Hymenobacter* (−1.21), and *Deinococcus* (−1.64), indicating taxa that were strongly depleted relative to other surfaces.

Together, these patterns demonstrated that Soil and Touchpoints harbour the most compositionally distinct microbial communities, characterised by broader, multi-taxon shifts rather than single dominant indicator genera. By contrast, the remaining surface types exhibited comparatively similar genera profiles with fewer pronounced deviations from the community-wide mean.

### Alpha and beta diversity highlight surface-specific microbial community patterns

Median alpha diversity varied only modestly across the five outdoor surface types (**Figure 2A**). Median Shannon diversity values were broadly similar for Pathways (3.01), Waterside (3.03), Touchpoints (2.99), and Soil (2.93), with no significant pairwise differences among these surfaces (Wilcoxon *p* > 0.05). Refuse surfaces showed the lowest median Shannon diversity (2.50) and were significantly less diverse than Pathways (*p* = 0.038) and Watersides (*p* = 0.018). No other comparisons reached significance. Additional alpha diversity metrics showed comparable but not identical patterns (**Supplementary Figure 6**). Median observed richness was highest on Touchpoints (56) and lowest in Soil (30), with Refuse samples at the middle of this range (53). Faith’s phylogenetic diversity medians followed a similar trend, ranging from 8.3 in Soil to 15.5 on Touchpoints, with Refuse again showing intermediate values (13.5). This indicated that although Refuse surfaces exhibited the lowest Shannon diversity, reductions were not consistently observed across all alpha diversity metrics.

**Figure 2.**
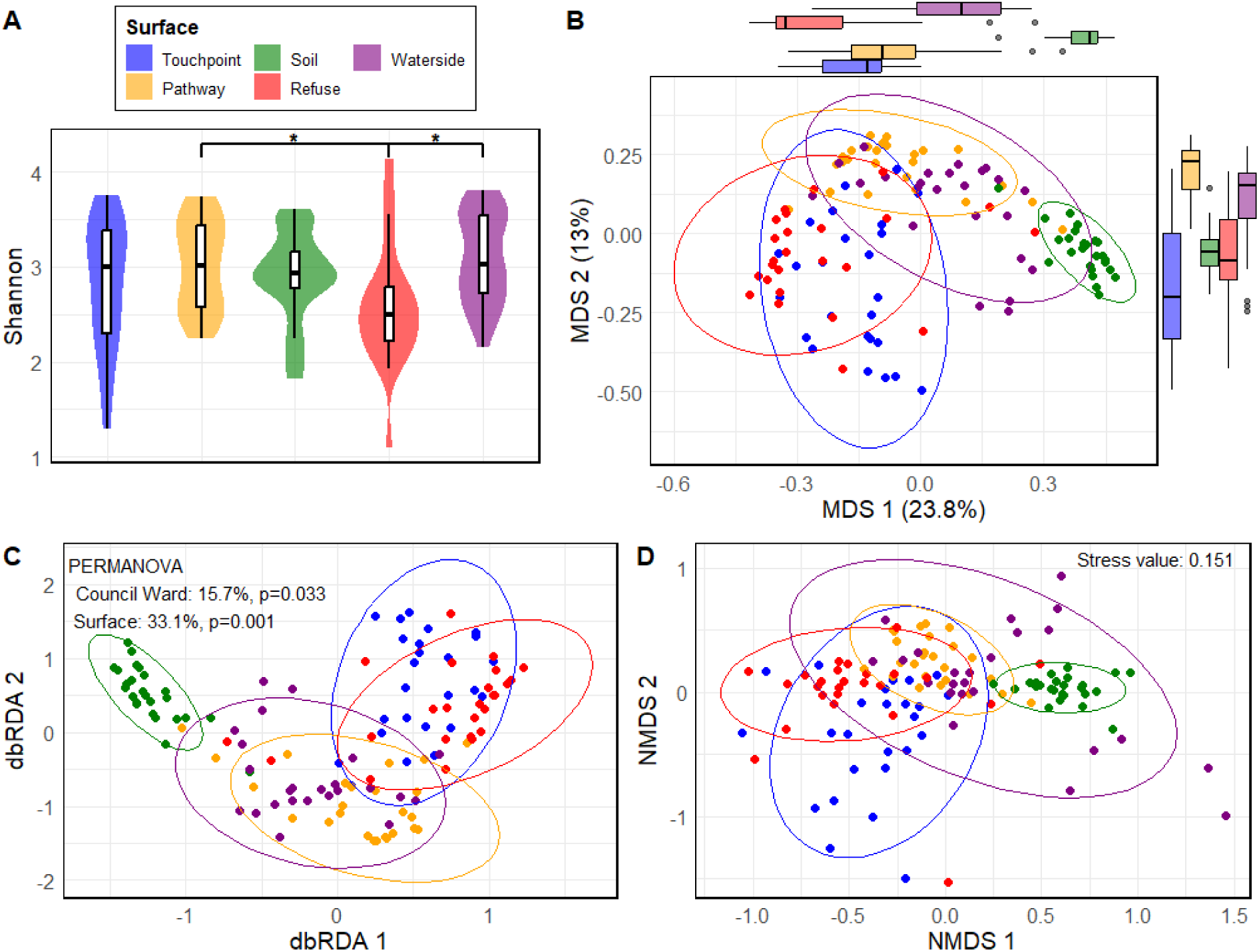
Alpha and beta diversity of microbiomes across the five outdoor surface types. **A)** Shannon diversity values, with asterisks indicating significant pairwise differences based on Wilcoxon tests (p < 0.05). **B–D)** Ordination of Bray–Curtis community dissimilarities visualised using **B)** unsupervised Multi-Dimensional Scaling (MDS), **C)** supervised distance-based Redundancy Analysis (dbRDA) constrained by Council Ward and Surface (PERMANOVA: Council Ward = 15.7%, p = 0.033; Surface = 33.1%, p = 0.001), and **D)** unsupervised non-metric MDS (NMDS) with an associated stress value of 0.151. Ellipses depict 95% confidence intervals around group centroids.

Beta diversity analyses showed clear differences in community composition among the five outdoor surface types (**Figure 2B**). Unsupervised Multi-Dimensional Scaling (MDS) ordination of Bray-Curtis dissimilarities revealed distinct clustering of samples by Surface, with Soil forming the most separated group. By contrast, Touchpoint and Refuse communities overlapped closely, as did Pathways and Waterside samples, indicating greater compositional similarity among these surface pairs. The first two MDS axes explained 23.8% and 13.0% of the variation in community composition respectively, indicating that a substantial proportion (36.8%) of the dissimilarity structure was captured in the two dimensional ordinations. The marginal boxplots further illustrate the strong separation of Soil along MDS1, while the remaining surfaces occupied broadly overlapping regions on both axes.

Distance-based redundancy analysis (dbRDA) was used to visualise community variation attributable to Surface and Council Ward (**Figure 2C**). Soil samples formed a distinct, isolated cluster, whereas the remaining surfaces showed overlapping community compositions. Touchpoint and Refuse surfaces clustered tightly together, while Pathway and Waterside samples also displayed close overlap, indicating shared compositional structure within these environment pairs. The amount of variation explained by Surface and Council Ward was quantified using Permutational Multivariate Analysis of Variance (PERMANOVA) on Bray-Curtis dissimilarities (**Supplementary Material 4**). When tested individually, only Surface showed a significant effect on community composition, explaining 33.1% of the variance (F = 14.64, p = 0.001), whereas Council Ward explained 15.7% but was non-significant (F = 0.77, p = 0.998). However, in the combined model, Surface and Council Ward together explained 48.7% of the variation and both factors were statistically significant (Surface: F = 15.30, p = 0.001; Council Ward: F = 1.21, p = 0.033). This shift highlighted that the strong Surface effect dominated the overall structure, and once this was controlled for, a smaller but significant Council Ward-level signal became apparent.

Tests for homogeneity of multivariate dispersion (PERMDISP) indicated that dispersion differed significantly among surfaces (F = 8.17; p = 0.001), suggesting that part of the Surface effect reflected differences in within-group heterogeneity. By contrast, no dispersion differences were observed among council wards (F = 1.11; p = 0.344), supporting the robustness of the Council Ward effect observed in the combined PERMANOVA model.

Non-metric MDS (NMDS) mapped the multivariate Bray–Curtis dissimilarities onto two-dimensional space, producing a stable ordination with a stress value of 0.151. As stress values closer to 0 indicate better agreement between the ordination and the original distance matrix, this value reflects a good representation of the underlying community structure (**Figure 2D**). The overall clustering pattern mirrored the MDS ordination, with Soil samples forming the most distinct group and the other surfaces showing varying degrees of overlap. Although the geometric arrangement of groups differed slightly, reflecting the arbitrary orientation of NMDS axes, the relative relationships among surface types remained consistent, underscoring the robustness of the observed community structure.

### Touchpoints dominate surface-specific variation in antimicrobial resistance gene abundance and diversity

Across all samples, Touchpoints exhibited the highest median overall sequencing-derived ARG abundance, with reads per kilobase per million (RPKM) values exceeding those of all other outdoor environments when utilising thresholds of >1 read, ≥90% nucleotide identity and 60% gene coverage (**Figure 3A**). Pairwise Wilcoxon tests confirmed that Touchpoints carried significantly greater sequencing-derived ARG loads than Pathways (p = 0.0016), Soil (p = 0.0376) and Waterside samples (p = 0.0045), while the comparison with Refuse approached significance (p = 0.0694). By contrast, all other surfaces displayed uniformly low ARG abundances, with no significant differences detected among Pathways, Soil, Refuse and Waterside (p ≥ 0.07). These findings indicated that the ARG burden on outdoor urban surfaces is heavily influenced by direct human contact. Exploratory analyses with the same thresholds at 30% gene coverage yielded the same overall pattern and further increased the strength of statistical differences between Touchpoints and all other surface types (**Supplementary Figure 7A**). ARG abundance showed no consistent spatial pattern when samples were grouped by Council Ward, with variation largely driven by the distribution of Touchpoint samples rather than geographical location (**Supplementary Figure 8**).

**Figure 3.**
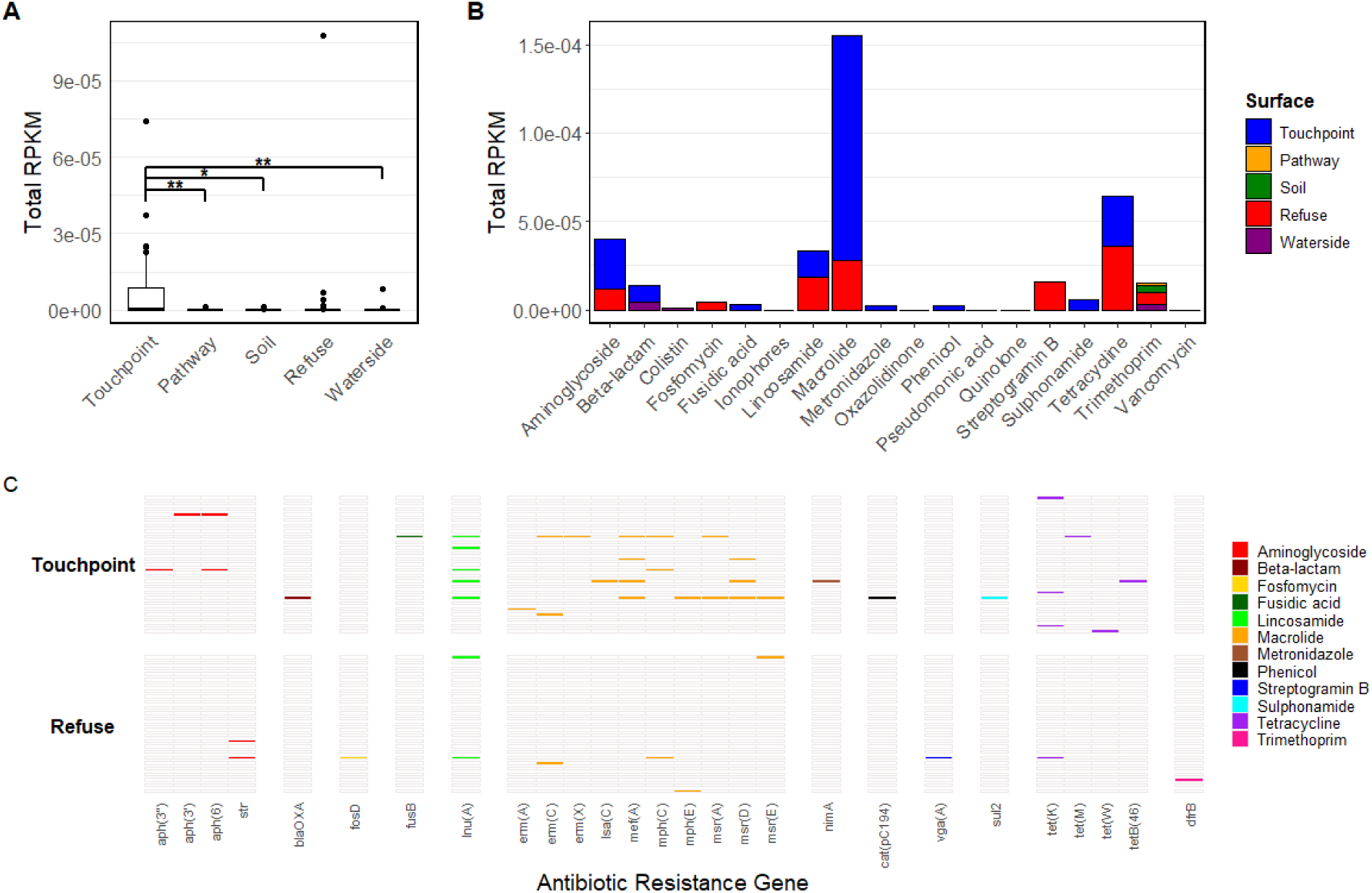
All panels are based on ARG hits meeting a ≥60% gene coverage threshold. **A)** Overall sequencing-derived antimicrobial resistance gene (ARG) abundance, calculated as the sum of RPKM-normalised ARG reads, across the five surfaces (Touchpoints, Pathways, Soil, Refuse and Waterside), with asterisks indicating significant pairwise differences based on Wilcoxon rank-sum tests (p < 0.05 = *, p < 0.01 = **). **B)** ARG abundance summed by antibiotic class for each surface type, expressed as RPKM. **C)** Presence-absence profiles of detected ARGs in Touchpoint and Refuse samples, grouped by antibiotic class.

To assess the potential influence of individual high-abundance observations on differences in overall ARG abundance, a sensitivity analysis was performed excluding the highest-RPKM observation from the Refuse group and the highest-RPKM observation from the Touchpoints group (**Supplementary Figure 9**). Following exclusion, the previously significant differences between Touchpoints and Pathways, Soil and Waterside remained significant, while the comparison between Touchpoints and Refuse also became significant (Wilcoxon rank-sum test, FDR-adjusted *p* = 0.0471). Thus, exclusion of the two highest-abundance observations did not alter the previously observed significant differences and resulted in a significant difference between Touchpoints and all other surface categories.

When ARGs were grouped by antibiotic class, Touchpoints again showed the broadest and highest abundance resistome profiles when utilising a 60% gene coverage threshold (**Figure 3B**). Macrolide-associated genes were the most abundant, followed by tetracycline, aminoglycoside and lincosamide determinants, all of which displayed elevated levels on Touchpoints. Notably, tetracycline- and lincosamide-associated genes also showed a higher proportional representation in Refuse samples, aligning with the limited yet detectable ARG signal previously observed for this surface type. Aside from Refuse, the remaining surface types displayed sparse ARG profiles, with most antibiotic classes either absent or present at very low abundance. These summed RPKM class-level patterns closely mirrored those identified at the surface level, indicating that the elevated resistome signal on Touchpoints reflected contributions from multiple antibiotic classes rather than a single dominant group. Analysis using a 30% gene coverage threshold revealed the same class-level trends, with Touchpoints again dominating the resistome signal; however, at this lower threshold, their proportional contribution increased further, mirroring the intensified differences observed at the surface level (**Supplementary Figure 7B**).

To examine the specific genes underlying the resistome patterns observed in Figures 3A and 3B, we assessed the presence–absence of all detected ARGs in Touchpoint and Refuse samples using a 60% gene coverage threshold (**Figure 3C**). Touchpoints displayed the broadest ARG repertoire, spanning multiple antibiotic classes, although most individual genes were detected only once or twice across the 25 samples. This indicated that while Touchpoints harboured a diverse set of ARGs, individual determinants generally occur at low frequency. Refuse samples contained a smaller but distinct subset of genes, also characterised by sparse and largely sample-specific detections.

Comparison with Figure 3B highlighted an important distinction between gene presence and summed RPKM values: despite Refuse having very few detectable tetracycline and lincosamide genes, single high-abundance detections produced relatively large total RPKM values for these classes. By contrast, Touchpoints showed lower per-gene abundance but a broader and more consistently detected repertoire across samples. This difference explained why Refuse appears more prominent for specific antibiotic classes in Figure 3B despite its low overall gene diversity.

Because ARG presence–absence at 60% coverage was highly sparse, we repeated the analysis at a 30% threshold (**Supplementary Figure 7C**). This recovered many additional low-abundance ARGs, especially in Touchpoint samples, yet the overall pattern remained unchanged: Touchpoints remained the dominant source of ARG diversity, followed by a much smaller contribution from Refuse. Thus, although higher coverage filtering reduced the number of detectable genes, the broader trends in ARG distribution were consistent across thresholds. It is important to note, however, that a 60% coverage threshold is not especially stringent, and lower coverage thresholds further increase the risk of false-positive calls, including truncated or non-functional gene fragments.

## Discussion

Surface identity was the strongest determinant of microbial community composition across the urban environments sampled in Liverpool, explaining 33.1% of variation in community structure. Soil and Touchpoint–Refuse communities showed distinct compositional profiles, while Pathway and Waterside communities occupied a more intermediate position within the broader urban microbiome. These differences were accompanied by marked variation in the distribution of ARGs, with Touchpoints and Refuse exhibiting the greatest resistome burden and Soil, Pathways and Waterside showing comparatively little ARG detection. Together, these findings suggest that urban surfaces represent distinct microbial microhabitats, shaped by differences in substrate, environmental exposure, moisture, and microbial sources, with implications for the distribution of AMR across the urban environment.

### Microbial Community Composition

Soil samples supported a compositionally distinct microbiome, consistent with the expectation that this surface would be shaped primarily by environmental rather than direct human-associated processes. Soil formed a distinct cluster in beta-diversity analyses and exhibited a taxonomic profile clearly differentiated from the other surface types. This distinction was evident at both phylum and genus levels, with Soil exhibiting the largest number of genera with strong positive or negative ALDEx2 effect sizes. Pseudomonadota, although abundant across all surface types, was significantly enriched in Soil (∼69%), while Acidobacteriota was also significantly more abundant than on other surfaces. Both phyla are widely associated with soil environments [32–36], supporting the interpretation that the observed differences reflect distinct soil-associated communities rather than simply variation in overall taxonomic richness.

The strong compositional separation of Soil was not accompanied by higher alpha diversity, indicating that its distinctiveness was driven primarily by community composition rather than greater taxonomic richness. Similar patterns have been reported in broader syntheses of urban microbiomes, which emphasised that even under intense anthropogenic influence, soil community structures remain distinct and heterogeneous [37, 38]. A global study of urban greenspace soils spanning 27 cities and 17 countries further identified consistent characteristics of urban soil microbiomes across geographically diverse environments [39]. These findings are broadly consistent with the distinct community structure observed in our urban Soil samples.

Touchpoints and Refuse appeared to represent a contrasting component of the urban microbial community, characterised by greater human and organic influence. Outside of phyla abundant across all surfaces, they were uniquely united by a significantly elevated abundance of Ascomycota, while Percolozoa were uniquely enriched on Touchpoints. Despite differences at the genus level, including high *Aureobasidium* enrichment on Refuse surfaces, beta-diversity analyses showed substantial overlap between Touchpoints and Refuse, indicating broadly similar overall community structures. Refuse also exhibited lower Shannon diversity despite intermediate richness- and phylogeny-based diversity, potentially reflecting greater dominance by a subset of taxa rather than reduced richness, as Shannon diversity is sensitive to evenness [40–42].

These findings are consistent with previous urban metagenomic studies demonstrating that microbial community composition varies according to surface type and human interaction. The global MetaSUB survey, for example, found that frequently contacted surfaces such as doorknobs, buttons and railings showed greater similarity to human skin microbiomes than less frequently contacted urban surfaces [43]. Similarly, studies of urban transit systems have reported enrichment of human-associated taxa on frequently contacted surfaces [44], while investigations of urban dustbin surfaces have identified contributions from both human-associated and environmental sources, with community composition varying according to waste type and sampling context [45].

Pathways and Watersides occupied a more intermediate position within the broader urban microbiome, with the two surface types showing substantial overlap across community-composition analyses. At the phylum level, both Pathways and Watersides were characterised by elevated relative abundances of Cyanobacteriota, which were significantly higher than on other surface types. In contrast to Soil and Touchpoints, they showed relatively little genus-level differentiation, while their communities formed strongly overlapping clusters in beta-diversity analyses. Their combined cluster also overlapped with the distributions of the other surface types, placing these environments more centrally within the overall urban microbiome. Similar intermediate and environmentally influenced community profiles have been reported in studies of exposed urban surfaces, including outdoor infrastructure at terrestrial–aquatic interfaces [46, 47].

### Methodological and Ecological Factors Shaping Microbial Communities

Interpretation of surface-specific differences should first consider the potential influence of sampling approach and substrate characteristics. Four of the five surface categories were sampled using the same swabbing procedure, providing consistency in collection across these environments. However, the physical properties of the substrates may have influenced the amount and composition of material recovered, as surfaces differ in characteristics such as texture, moisture and the accumulation of biological material. These differences could therefore contribute to variation between surface categories independently of underlying community composition.

Soil represents the clearest methodological distinction, as material was collected directly from the soil rather than by swabbing a defined surface area. The relatively tight clustering of Soil samples may therefore partly reflect the greater consistency of this collection approach compared with swabbing heterogeneous surfaces. Despite comparable sequencing depth, Soil had a markedly lower proportion of taxonomically classified reads, consistent with the underrepresentation of environmentally derived genomes in commonly used reference databases [48–52]. Together, these factors introduce some uncertainty into the interpretation of taxonomic differences between Soil and the other surfaces.

The distinctness of the Soil microbiome likely reflects the different physical and ecological conditions of this substrate, which provide microbial niches and sources that differ from those associated with built and human-contact surfaces. The samples captured the microbial community associated with the accessible soil surface, where communities may be shaped by local environmental conditions and inputs from surrounding vegetation, atmospheric deposition and human activity. This is consistent with evidence that urbanisation and associated environmental stressors can alter soil microbial communities, including through selection for disturbance-tolerant taxa [53–56]. The distinctness of the Soil microbiome therefore likely reflects the different physical and ecological conditions of this substrate, which provide microbial niches and sources that differ from those associated with built and human-contact surfaces.

Touchpoints were explicitly associated with human interaction, while Refuse was indirectly exposed to human contact through waste disposal and handling. Both surface types were uniquely enriched in Ascomycota, which are commonly associated with the human skin mycobiome and airborne and dust-associated microbial communities [57, 58], while also being important decomposers of organic waste [59]. This shared enrichment is therefore consistent with contributions from both human-associated and organic environmental sources. Many Touchpoint-enriched genera were established human-associated taxa [15, 60–63], consistent with repeated direct contact, whereas Refuse showed fewer strongly differentiated genera, suggesting a more heterogeneous assemblage influenced by mixed environmental and anthropogenic inputs.

The Refuse samples were predominantly collected from the external surfaces of litter and wheelie bins rather than directly from the waste contained within them. Thus, although some material associated with waste may have been deposited on these surfaces, visibly mouldy or decaying waste was not specifically targeted for sampling. The strong enrichment of *Aureobasidium* on Refuse, a genus widely associated with decaying organic material and plant and built-environment surfaces [64], may therefore reflect a combination of exposure to organic material and environmental deposition alongside human contact, rather than direct sampling of decomposing waste. This mixed exposure may help explain the combination of fungal and human-associated taxa observed on Refuse surfaces.

Percolozoa were uniquely enriched on Touchpoints, providing a further indication that local microhabitat conditions may contribute to community differences between surface types. Members of this group are widely distributed in moist soils, freshwater and biofilm communities where intermittent moisture and organic matter support their persistence [65], and free-living amoebae have also been reported in engineered water systems and drinking-water distribution networks [66, 67]. Their enrichment on Touchpoints may therefore reflect opportunistic colonisation of moist micro-niches associated with rain splash, condensation or biofilm formation on frequently touched infrastructure. Further targeted studies would be needed to establish the drivers and prevalence of Percolozoa on human-contact surfaces in urban environments.

Pathways and Watersides were more environmentally exposed than the human-associated surfaces, with Waterside samples collected from the edges of waterbodies and drainage channels where microbial communities are likely influenced by surface runoff and inputs from surrounding environments. These samples therefore represent environmentally exposed runoff zones rather than deeper aquatic or sewage-associated microbiomes. The relatively high abundance of Cyanobacteriota in Pathways (24%) and Watersides (12%) is consistent with the environmental exposure and moisture-associated conditions of these surfaces, as Cyanobacteriota are widely distributed across terrestrial and aquatic environments and can persist on sun-exposed and moisture-associated outdoor surfaces [68–70]. More broadly, continual microbial inputs from surrounding air, soil and water, including atmospheric deposition and other dispersal processes, may contribute to the relatively mixed communities observed on these surfaces [71, 72].

### Antimicrobial Resistance Gene Presence

Touchpoints were the principal resistome-bearing surfaces in our dataset, with Touchpoints exhibiting the highest and most consistently elevated ARG abundance across surface types. Given that these surfaces were selected to capture frequent human contact, the enrichment of ARGs is consistent with a contribution from human-associated microbial deposition. This aligns with metagenomic studies of public transport systems and indoor built environments, where high-touch surfaces have been associated with elevated resistome signals [15, 19, 43, 73–76]. In our dataset, Touchpoints also showed enrichment of human-associated bacterial genera including *Staphylococcus*, *Corynebacterium*, and *Streptococcus*, alongside resistance genes spanning multiple clinically relevant classes. These genera are recognised members of the human microbiome and can harbour diverse resistance mechanisms, including the macrolide-lincosamide-streptogramin, tetracycline and aminoglycoside resistance genes detected in our samples [77, 78]. Together, these findings are consistent with human contact contributing both bacteria and ARGs to outdoor infrastructure, although further work would be required to distinguish repeated deposition from longer-term persistence or colonisation.

Refuse also showed elevated ARG abundance, but this pattern was strongly influenced by a single high-abundance sample, which accounted for the majority of detected ARGs and RPKM within the surface category. Excluding this observation, together with the highest-RPKM Touchpoint sample, resulted in Touchpoints becoming significantly higher than Refuse as well as the other surface categories. Although the Refuse observation cannot be assumed to represent an outlier, its influence highlights the heterogeneity of the resistome associated with these surfaces. The Refuse samples were predominantly collected from the external surfaces of litter and wheelie bins, which may be exposed to variable combinations of human contact and waste-associated material. This heterogeneous exposure may therefore contribute to the variable ARG abundance observed across Refuse samples.

In contrast, ARGs were rarely detected in Pathways and Watersides and were exceptionally low in Soil. The absence or low abundance of detected ARGs in these environments should not necessarily be interpreted as an absence of resistance determinants, particularly given the recognised underrepresentation of environmental ARGs in reference databases [79, 80]. Soil is considered an important reservoir of both known and novel resistance determinants, while detectable ARG abundance can vary with environmental conditions and selective pressures [81, 82]. Similarly, outdoor Pathway and Waterside surfaces are exposed to UV radiation, desiccation and nutrient limitation, which may constrain the persistence of deposited ARG-bearing bacteria [83–85]. Overall, the resistome patterns suggest a greater association of detectable ARGs with surfaces exposed to human activity and waste-associated inputs than with more environmentally exposed surfaces.

ARG abundance was estimated from metagenomic sequencing data and normalised by RPKM rather than quantified directly using qPCR or ddPCR. The ARG abundance reported here therefore represents a sequencing-derived relative measure and should not be interpreted as an absolute ARG burden or copy number. Absolute quantification, potentially alongside an appropriate measure of microbial biomass, would provide complementary information on the quantity of resistance determinants present across surface types.

### Study Limitations

A key limitation of these data is the combined constraint of sequencing depth (∼12 million reads per sample; ∼2 Gb per metagenome) and the inherently low microbial biomass associated with outdoor surface swab samples. In such systems, limited DNA input and the presence of substantial non-target/background DNA reduce the effective coverage of individual microbial genomes, thereby constraining both detection sensitivity and downstream analytical resolution. Notably, parallel processing by an alternative provider failed to generate sufficient DNA for sequencing, further indicating that low biomass and DNA recovery were fundamental limitations and likely constrained downstream analyses.

Two low-biomass negative controls were included during sample processing: a blank collection swab and a blank DNA/RNA Shield buffer control. Both controls yielded 0 ng/µL DNA following extraction, while the corresponding library preparations yielded 0 and 0.01 ng/µL, respectively. However, sequencing data for these controls were not retained, preventing assessment of their taxonomic and resistome profiles. Although the very low DNA and library yields provide no indication of substantial contamination during sample processing, low-level contamination cannot be completely excluded in the absence of the corresponding sequencing data.

As a result of sequencing depth, even on Touchpoints where ARG diversity was relatively elevated, individual ARGs were often sparsely detected, with many genes present in only one or two samples. This limited statistical power for gene-level inference and necessitated interpretation at the level of surface-wide trends rather than individual ARGs. Similarly, the limited effective coverage constrained taxonomic resolution, and community composition was therefore assessed at the genus level rather than relying on species-level assignments, which were less consistently supported by the available sequencing data.

These constraints also impacted assembly-based approaches. Attempts to recover metagenome-assembled genomes (MAGs) and to assign ARGs to host backgrounds did not yield robust outputs, likely reflecting insufficient effective coverage per organism rather than sequencing depth alone. Consequently, detailed assessment of ARG mobility potential and horizontal gene transfer could not be performed with confidence.

While increased sequencing depth may partially mitigate these limitations, improved biomass recovery (e.g., through sampling strategy or DNA input optimisation) would likely be required to enable reliable genome-resolved and mobility-focused analyses in this type of low-biomass urban environment.

Importantly, however, the overarching pattern was robust: Touchpoints consistently exhibited a significantly higher overall sequencing-derived ARG abundance than other surface types. This signal was further strengthened when the gene coverage threshold was relaxed from 60% to 30%, increasing detection sensitivity at the expense of stricter gene-level confidence, while preserving the relative ordering of surfaces.

## Conclusions

We presented a city-wide metagenomic survey of outdoor public surfaces, providing a baseline characterisation of microbial community structure, diversity, and ARG distribution across distinct surface types. Our findings demonstrated that surface function and patterns of human interaction are key drivers of microbial assembly in outdoor urban environments. Touchpoints were consistently enriched for human-associated taxa and carried the highest resistome burdens, whereas Pathways and Watersides represented more transitional microbial environments with lower ARG prevalence. Although individual ARG detections were sparse, the surface-level patterns were robust, highlighting the importance of considering ecological context and human behaviour when interpreting resistome signals in the built environment. Together, these results emphasised the value of outdoor metagenomic surveillance for understanding how urban infrastructure and human activity shape microbial and resistance landscapes, and provide a framework for future studies assessing temporal dynamics, environmental persistence, and potential public health relevance.

## Author Contributions

G.A.J. was the lead author and A.P.R. supervised the study. G.A.J. and A.P.R. conceptualised the study. G.A.J., A.M.O., A.H., E.A., P.A., A.B., A.C., K.D., R.G., A.K., A.M., S.M., R.P., M.B.R., P.S., E.S., K.C., M.M., H.M. and A.P.R. contributed to methodology development. G.A.J., A.M.O., P.A., A.B., A.C., K.D., R.G., A.K., A.M., S.M. and P.S. conducted sample collection. A.H. and H.M. conducted sample processing and genome sequencing. G.A.J. performed formal data analysis and wrote the original draft manuscript. G.A.J., A.M.O., A.H., E.A., P.A., A.B., A.C., K.D., R.G., A.K., A.M., S.M., R.P., M.B.R., P.S., E.S., K.C., M.M., H.M. and A.P.R. contributed to manuscript review and editing.

