## Supplementary Figures for "A citywide metagenomic analysis reveals surface-specific microbiome and resistome patterns in outdoor urban environments across Liverpool, UK"

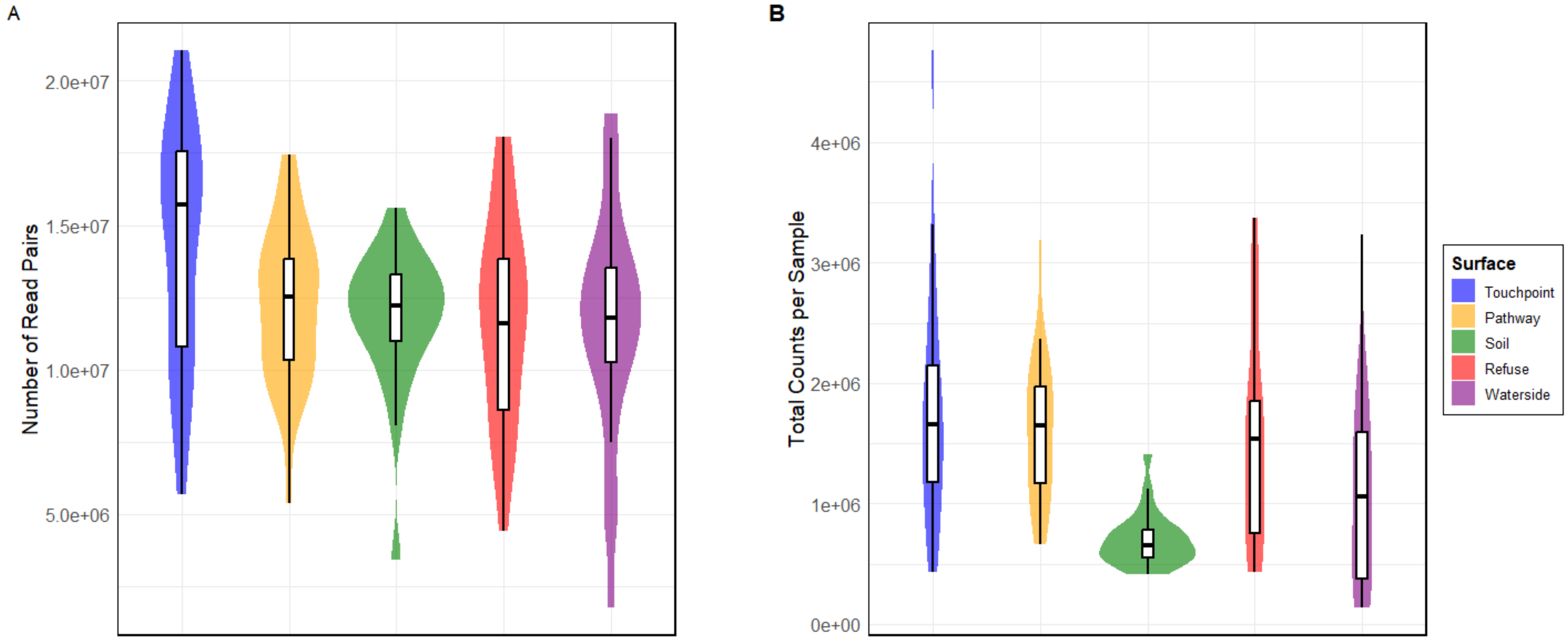


**Supplementary Figure 1.** Total (A) number of read pairs and (B) number of reads mapped to taxa per sample across each surface type. Violin plots represent the distribution of values, while overlaid boxplots show the interquartile range (25th-75th percentile) and median; whiskers extend to 1.5× the interquartile range.

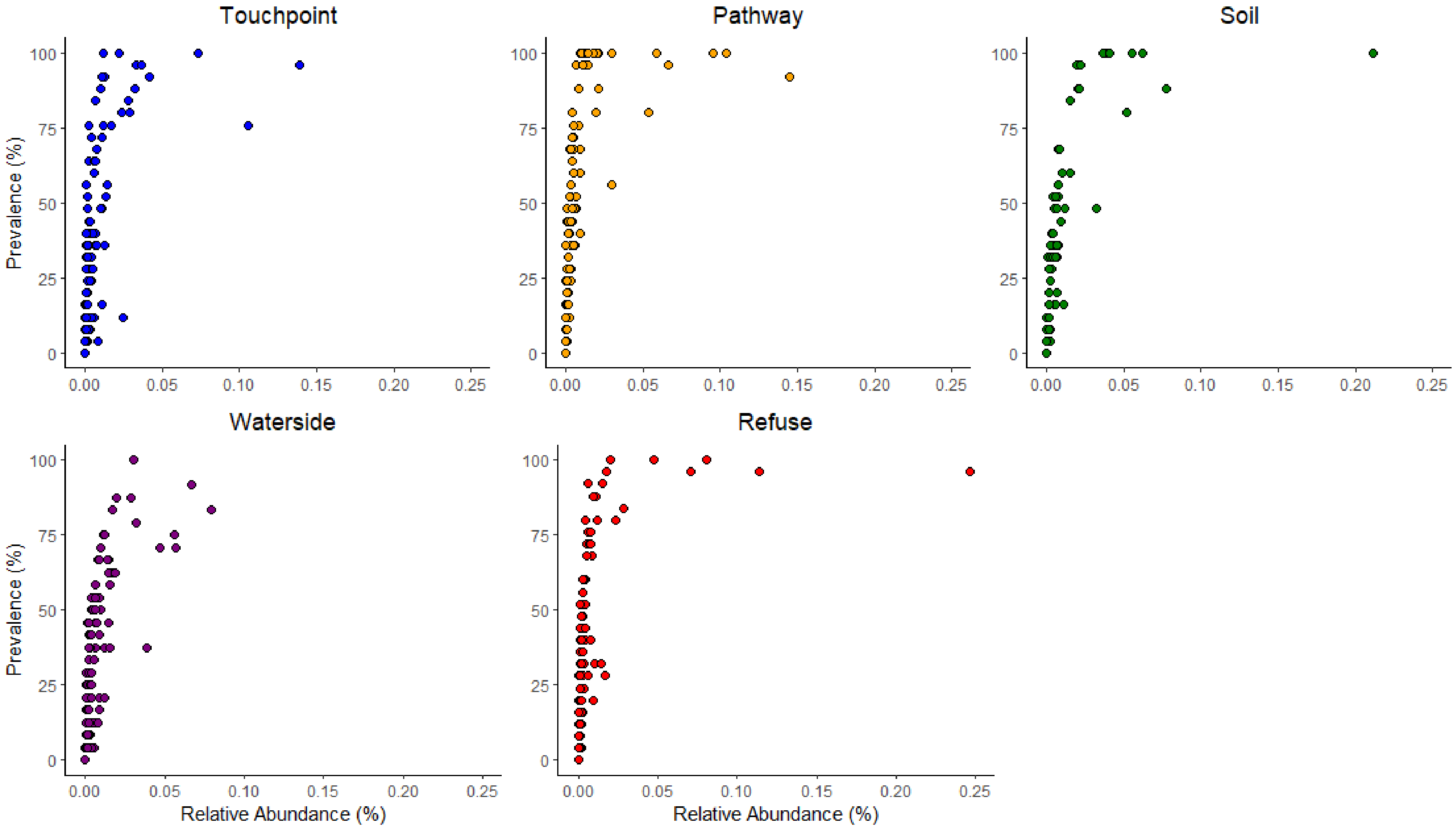


**Supplementary Figure 2.** Relative abundance (%) vs prevalence (%) of unique bacterial genera detected on each surface type. Each point represents a genus, where relative abundance refers to the proportion of sequencing reads assigned to that genus, and prevalence denotes the percentage of samples within each surface type in which the genus was detected.

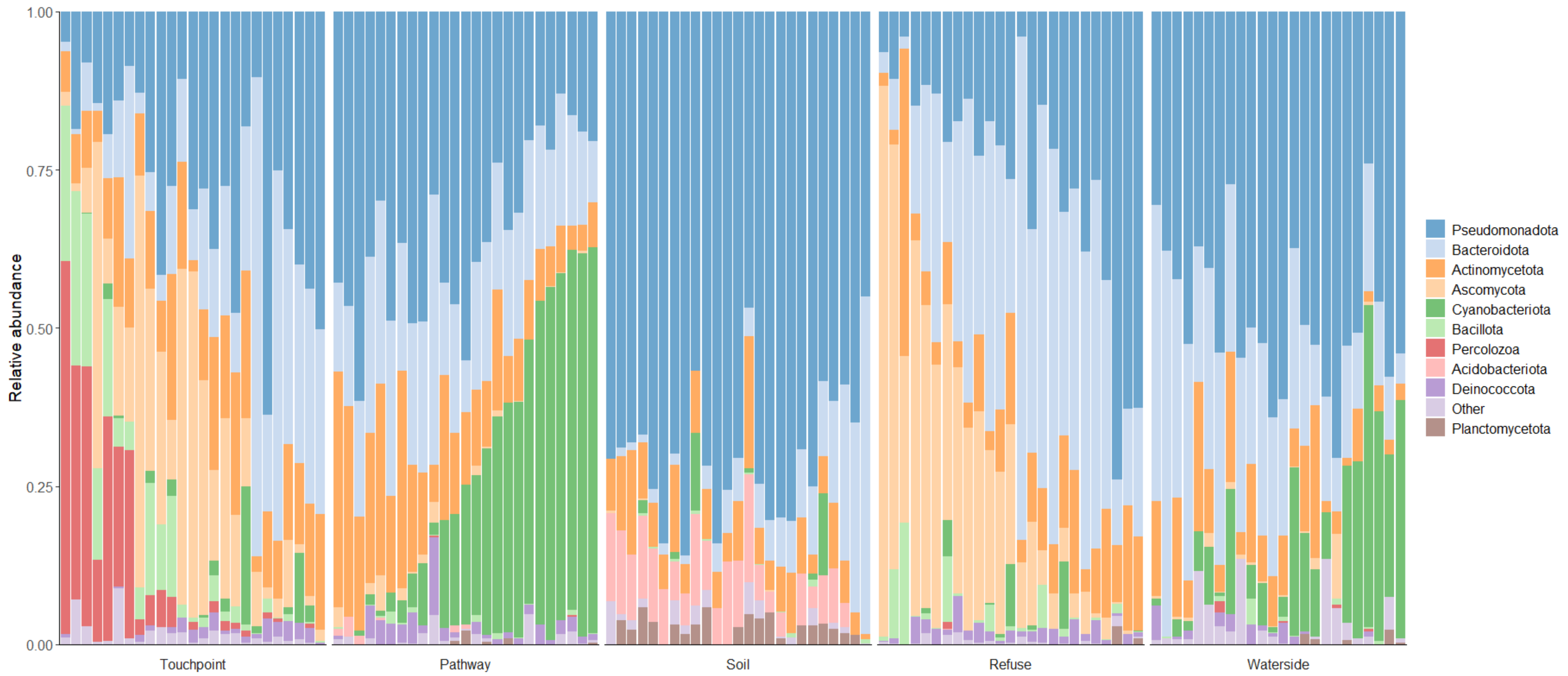


**Supplementary Figure 3**. Per-sample phylum-level taxonomic composition across surface categories. Each stacked bar represents one sample, and colours indicate the ten most abundant phyla identified across the full dataset; all remaining phyla are combined as “Other”. Bar heights show relative abundance. Within each surface, samples are ordered according to their score on the first principal-coordinate axis derived from Bray–Curtis dissimilarities of phylum-level relative-abundance profiles.

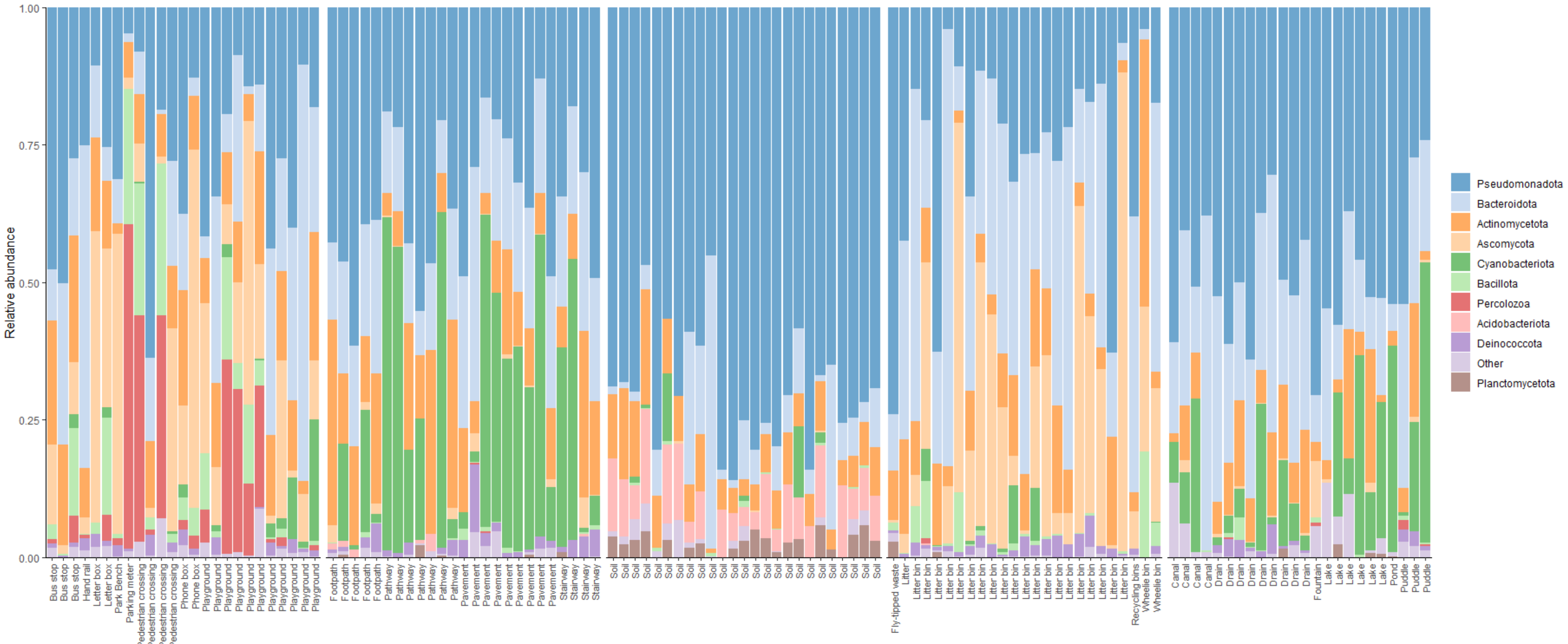


**Supplementary Figure 4**. Per-sample phylum-level taxonomic composition across surface categories, ordered by surface subtype metadata. Each stacked bar represents one sample, and colours indicate the ten most abundant phyla identified across the full dataset; all remaining phyla are combined as "Other". Bar heights show relative abundance. Within each surface, samples are ordered alphabetically by their surface subtype metadata (for example, drain, playground, or litter bin).

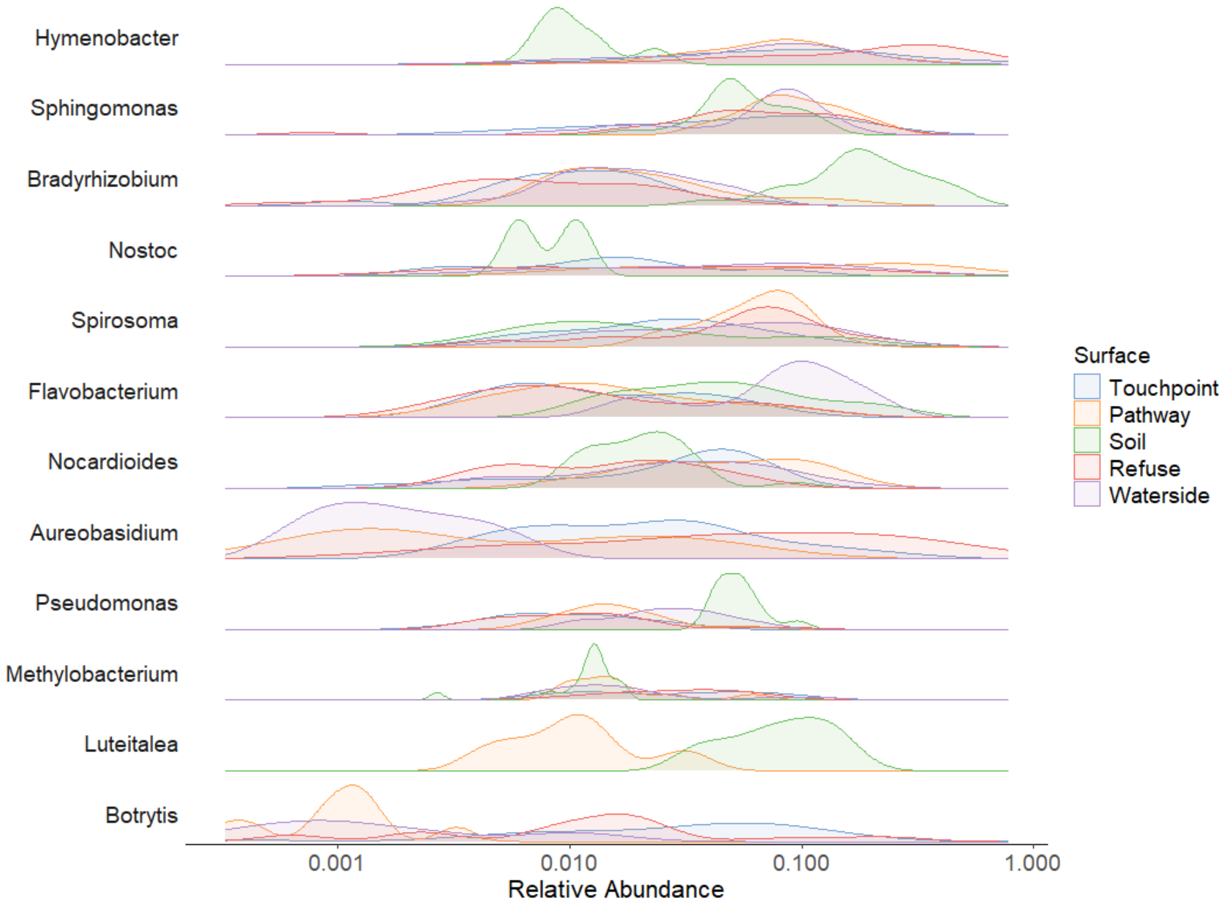


**Supplementary Figure 5**. Density plots showing the relative abundance of genera across surface types. The five most abundant genera within each surface type were selected and merged across groups, resulting in a final set of twelve unique genera.

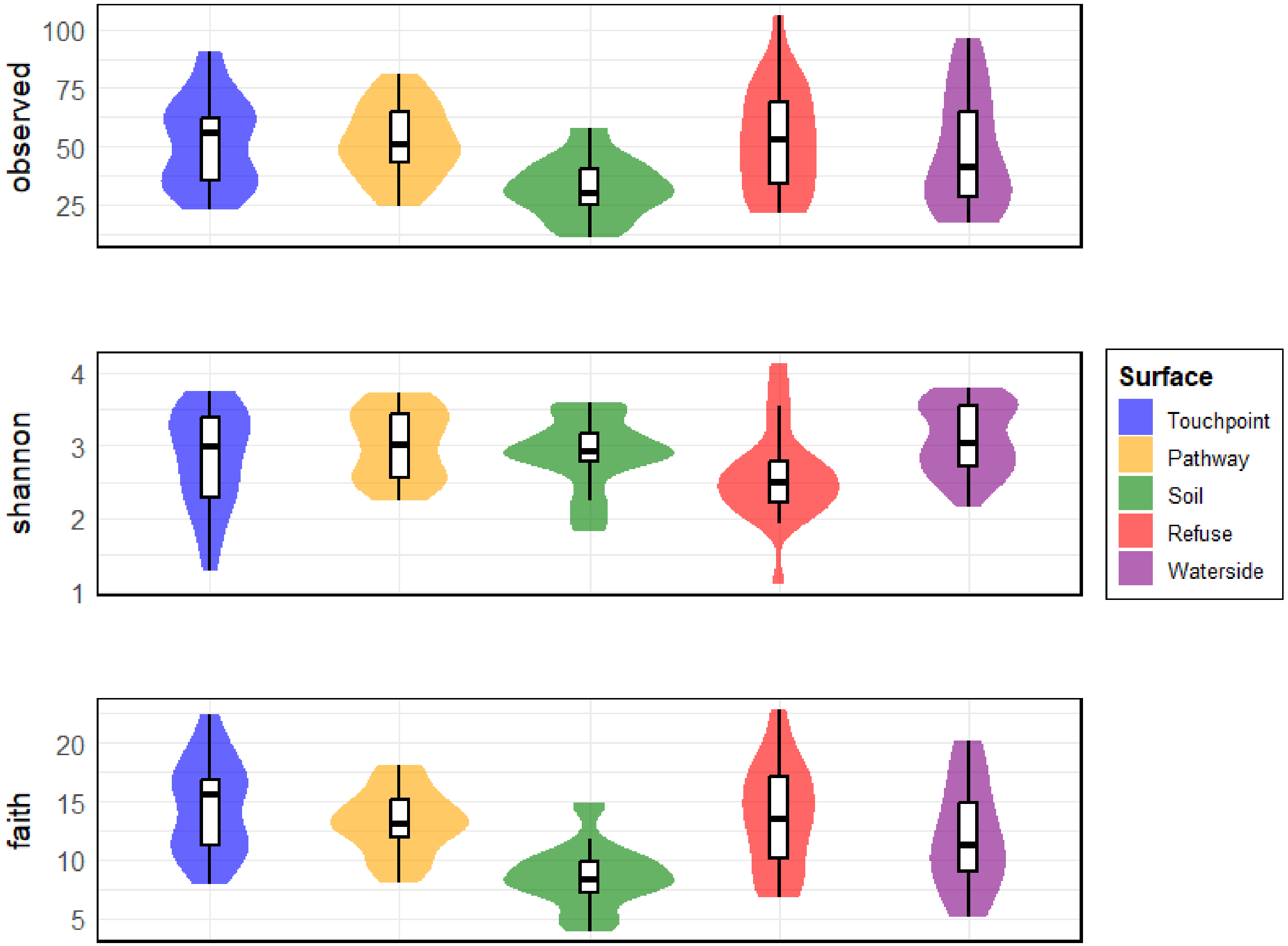


**Supplementary Figure 6**. Observed richness, Shannon diversity, and Faith's phylogenetic diversity across the five outdoor surface types.

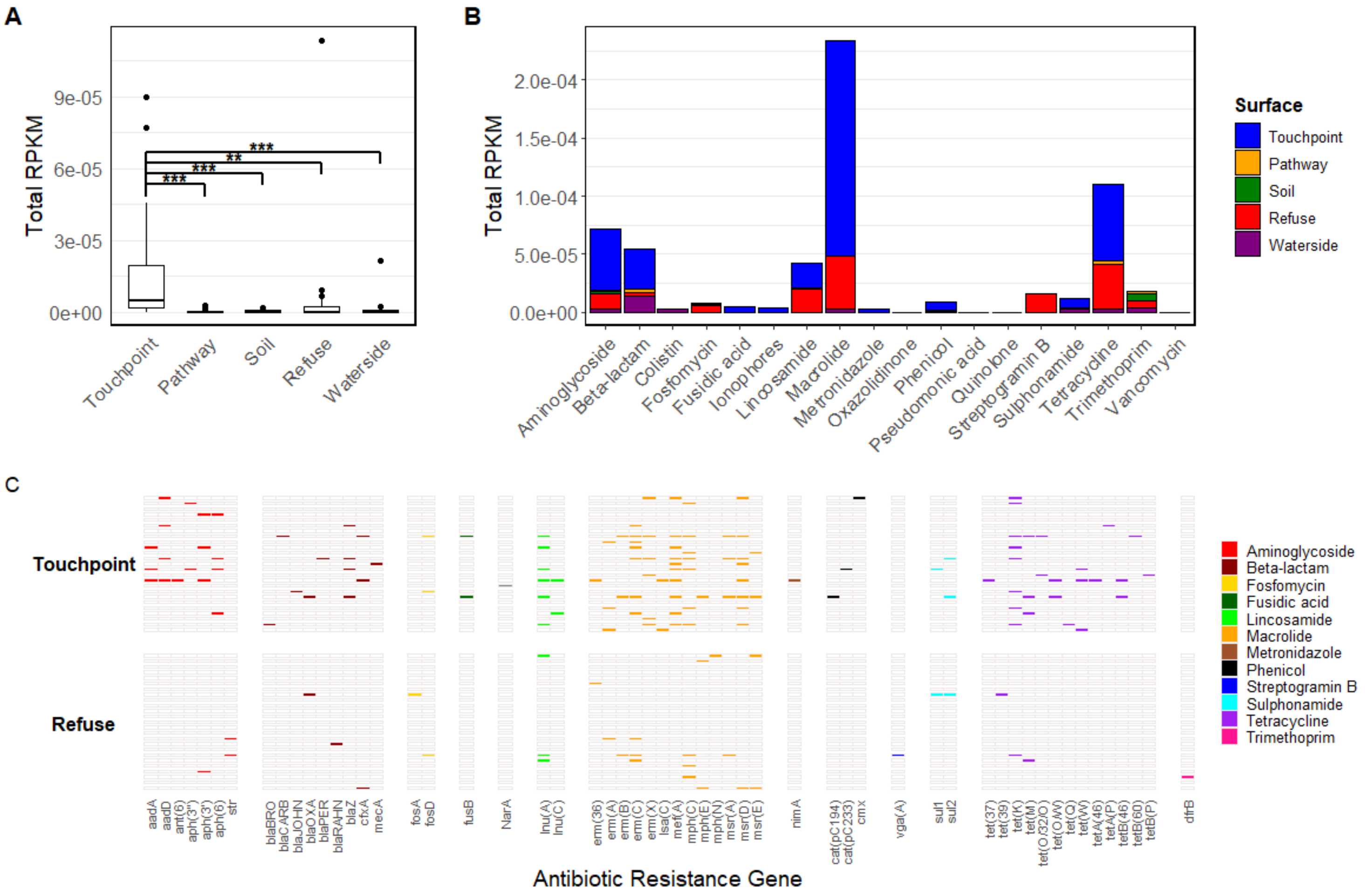


**Supplementary Figure 7.** All panels are based on ARG hits meeting a ≥30% gene coverage threshold. **A)** Overall sequencing-derived antimicrobial resistance gene (ARG) abundance, calculated as the sum of RPKM-normalised ARG reads, across the five surfaces (Touchpoints, Pathways, Soil, Refuse and Waterside), with asterisks indicating significant pairwise differences based on Wilcoxon rank-sum tests ($p < 0.05$ = *, $p < 0.01$ = **, $p < 0.001$ = ***). **B)** ARG abundance summed by antibiotic class for each surface type, expressed as RPKM. **C)** Presence-absence profiles of detected ARGs in Touchpoint and Refuse samples, grouped by antibiotic class.

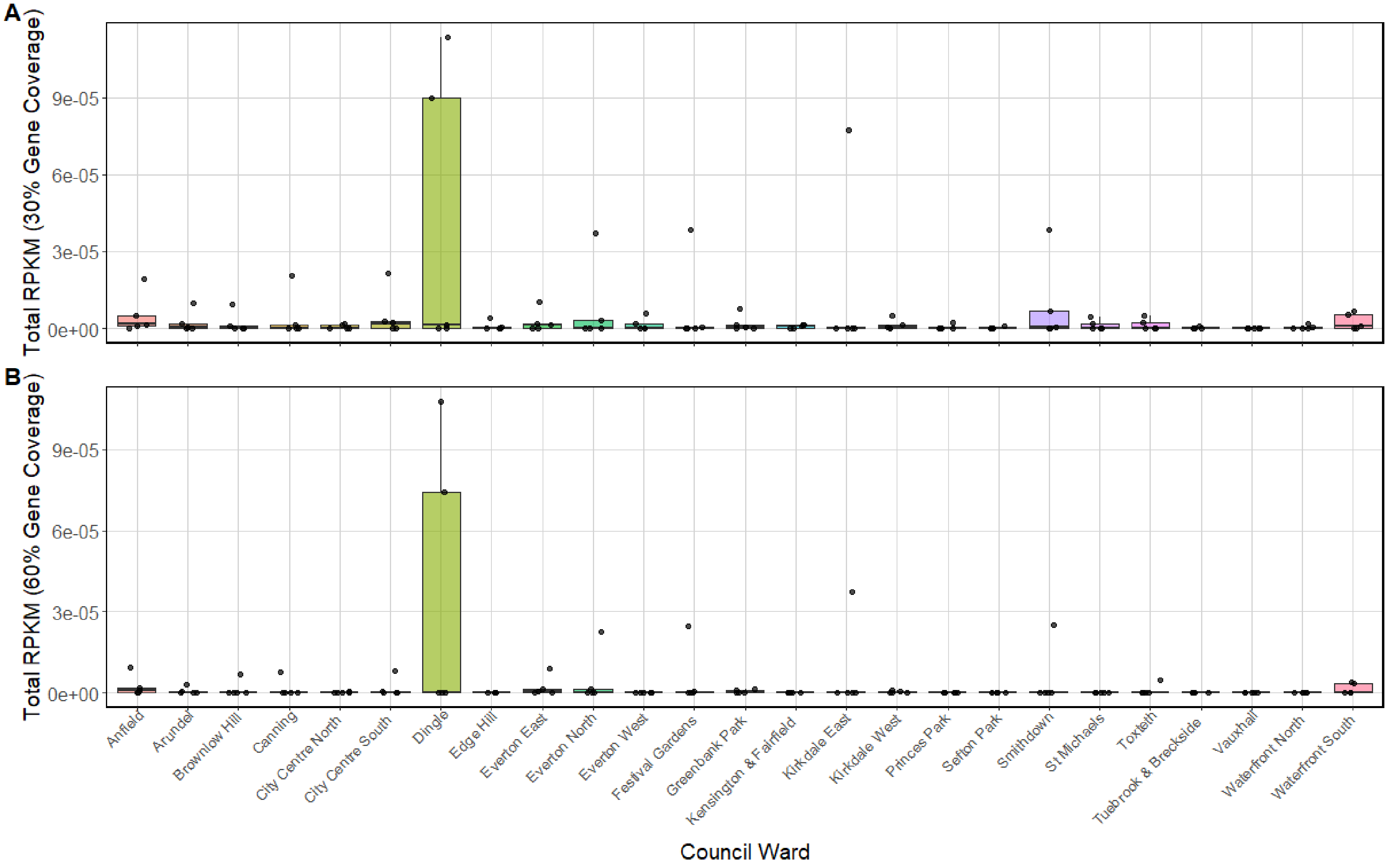


**Supplementary Figure 8.** Total antimicrobial resistance gene (ARG) abundance (reads per kilobase per million, RPKM) across the 25 Council Wards for **A)** 30% and **B)** 60% gene coverage thresholds. Each point represents an individual sample; boxplots show the distribution within each ward.

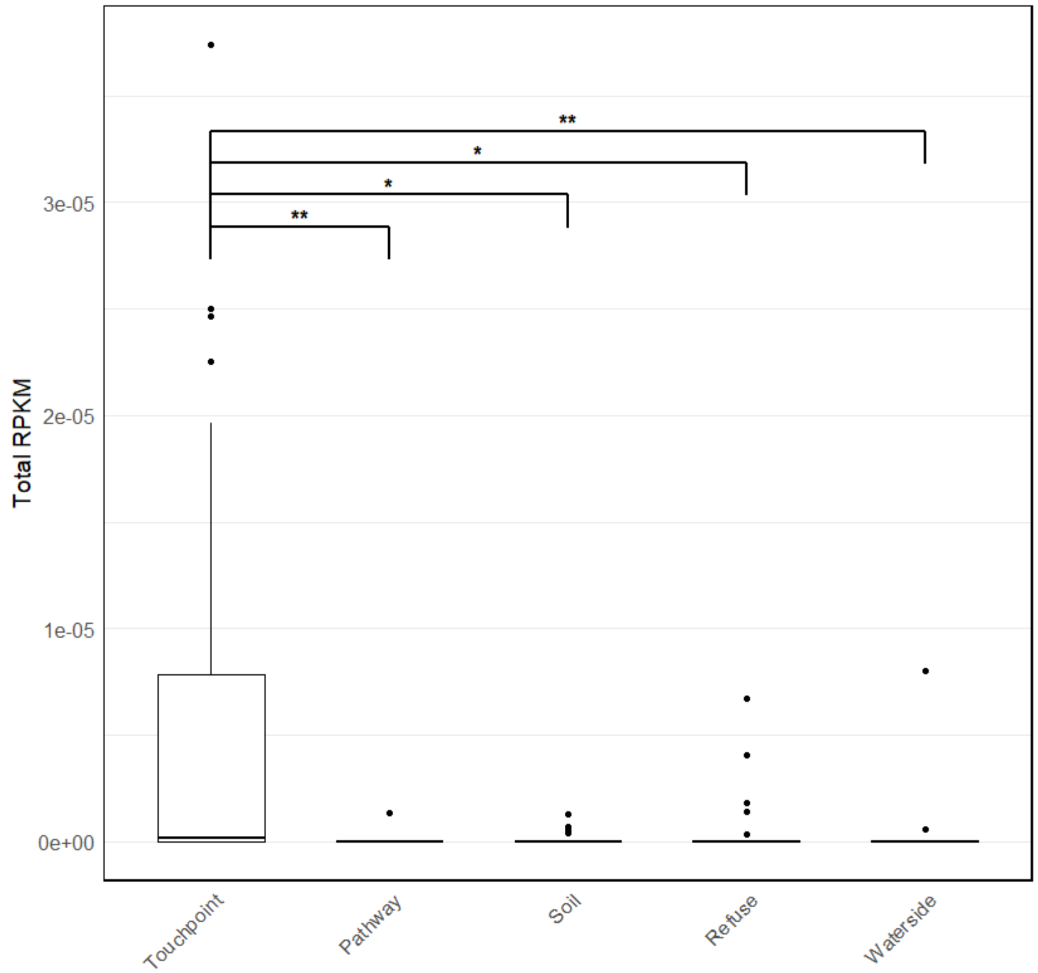


**Supplementary Figure 9**. Sensitivity analysis of overall sequencing-derived antimicrobial resistance gene (ARG) abundance following exclusion of the highest-RPKM observations. Overall sequencing-derived ARG abundance, calculated as the sum of RPKM-normalised ARG reads, across the five surface categories (Touchpoints, Pathways, Soil, Refuse and Waterside) after excluding the highest-RPKM observation from the Refuse group and the highest-RPKM observation from the Touchpoints group. ARG hits meeting a ≥60% gene coverage threshold were included. Asterisks indicate significant pairwise differences based on Wilcoxon rank-sum tests ($p < 0.05$ = *, $p < 0.01$ = **).
